# ATF4 Dependent Increase in Mitochondrial-Endoplasmic Reticulum Tethering Following OPA1 Deletion in Skeletal Muscle

**DOI:** 10.1101/2022.09.12.507669

**Authors:** Antentor Hinton, Prasanna Katti, Margaret Mungai, Duane D. Hall, Olha Koval, Jianqiang Shao, Zer Vue, Edgar Garza Lopez, Rahmati Rostami, Kit Neikirk, Jessica Ponce, Jennifer Streeter, Brandon Schickling, Serif Bacevac, Chad Grueter, Andrea Marshall, Heather K. Beasley, Young Do Koo, Sue C. Bodine, Nayeli G. Reyes Nava, Anita M. Quintana, Long-Sheng Song, Isabella Grumbach, Renata O. Pereira, Brian Glancy, E. Dale Abel

## Abstract

Mitochondria and endoplasmic reticulum (ER) contact sites (MERCs) are protein- and lipid-enriched hubs that mediate interorganellar communication by contributing to the dynamic transfer of Ca^2+^, lipid, and other metabolites between these organelles. Defective MERCs are associated with cellular oxidative stress, neurodegenerative disease, and cardiac and skeletal muscle pathology via mechanisms that are poorly understood. We previously demonstrated that skeletal muscle-specific knockdown (KD) of the mitochondrial fusion mediator optic atrophy 1 (OPA1) induced ER stress and correlated with an induction of Mitofusin-2, a known MERC protein. In the present study, we tested the hypothesis that *Opa1* downregulation in skeletal muscle cells alters MERC formation by evaluating multiple myocyte systems, including from mice and *Drosophila*, and in primary myotubes. Our results revealed that OPA1 deficiency induced tighter and more frequent MERCs in concert with a greater abundance of MERC proteins involved in calcium exchange. Additionally, loss of OPA1 increased the expression of activating transcription factor 4 (ATF4), an integrated stress response (ISR) pathway effector. Reducing *Atf4* expression prevented the OPA1-loss-induced tightening of MERC structures. OPA1 reduction was associated with decreased mitochondrial and sarcoplasmic reticulum, a specialized form of ER, calcium, which was reversed following ATF4 repression. These data suggest that mitochondrial stress, induced by OPA1 deficiency, regulates skeletal muscle MERC formation in an ATF4-dependent manner.

## Introduction

The endoplasmic reticulum (ER) and mitochondria continuously exchange molecules to sustain intracellular homeostasis. This is facilitated, in part, by physical contacts between the two organelles known as mitochondria-associated membranes (MAMs) or mitochondrial-ER contact sites (MERCs) (Rieusset, 2018). MERC dysfunction, including alterations in MERC formation and morphology, has been associated with disease states characterized by altered mitochondrial oxidative metabolism and insulin resistance including obesity, type II diabetes (T2D), cardiovascular disease and aging (Rieusset, 2018).

MERC microdomains are enriched in proteins involved in mitochondrial Ca^2+^ flux, lipid transfer, and the regulation of mitochondrial function (Tubbs et al., 2014, 2018). The size, density, and thickness of MERCs regulate their function, and changes in MERC distances can modulate Ca^2+^ exchange, lipid homeostasis, and cell death (Csordás et al., 2018; Delprat et al., 2019; Giacomello & Pellegrini, 2016; Hirabayashi et al., 2017; Rieusset, 2018).

Several important MERC-tethering proteins are expressed in the MERC microdomain, including Mitofusin-2 (MFN2), the inositol 1,4,5-triphosphate receptor (IP_3_R3), glucose-related protein 75 (GRP75), voltage-dependent anion channel (VDAC) complex, the vesicle-associated membrane proteins B and C, and others. Most known MERC proteins are located on the outer mitochondrial membrane (OMM). A relationship between the inner mitochondria membrane protein optic atrophy 1 (OPA1) and regulation of MERCs, was recently described (Cartes-Saavedra et al., 2022). However, a complete understanding of mechanisms linking inner mitochondrial membrane proteins, such as OPA1, to the regulation of MERC formation remains to be achieved.

We previously showed that *OPA1* knockdown (KD) in skeletal muscle (OPA1smko/KD) impairs mitochondrial fusion and bioenergetics, induces ER stress, and correlates with the induction of MFN2 (Pereira et al., 2017; Tezze et al., 2017). *OPA1* KD in skeletal muscle also increases activating transcription factor (ATF4) levels (Pereira et al., 2017). Interestingly, ATF4 activation is involved in several stress processes, including having a key role in the integrated stress response (ISR) (Costa-Mattioli & Walter, 2020; Hartwick Bjorkman & Oliveira Pereira, 2021; Kasai et al., 2019; Sasaki et al., 2020). ATF4 is also linked to muscle atrophy (Pereira et al., 2017). Skeletal muscle atrophy and reduced exercise activity are associated with MERC changes (Romanello & Sandri, 2015). Age-related physical inactivity is also associated with muscle loss and reduced OPA1 levels (Tezze et al., 2017). However, the mechanisms by which ATF4 may influence the morphology of MERC microdomains in these and other contexts are incompletely understood.

We hypothesized that OPA1-induced mitochondrial stress increases MERC abundance and narrows MERC distance, potentially as a response to ER stress. To test this hypothesis, we used OPA1 loss of function in myotubes, mouse skeletal muscle, and *Drosophila* flight muscle to assess the effects of OPA1 deficiency on MERC structure. Across models, we demonstrate that OPA1 deficiency in muscle cells induces the expression of MERC proteins and increases MERC tethering. Additionally, we find that the MERC structural changes in OPA1-deficient muscle cells are mediated, at least in part, by ATF4. Indeed, *Atf4* loss reverses the narrowing of MERC distances driven by *Opa1* loss, while *Atf4* overexpression alone is sufficient to increase MERC length and narrow MERC distances. This increase in MERC formation may represent an initially compensatory, but ultimately maladaptive mechanism that attempts to restore calcium homeostasis in response to OPA1-deficiency-induced mitochondrial stress.

## Materials and Methods

### Animal Studies

The University of Iowa Institutional Animal Care and Use Committee (IACUC) approved all protocols for animal work. For primary satellite cells isolation, 16 male mice on the C57BL/6J genetic background were used. We previously reported that male and female mice demonstrate similar effects to loss of *OPA1* (Pereira et al., 2017). For all gastrocnemius samples, mice were aged to 9 months.

Generation of OPA1 KO animals and TUDCA treatment was achieved using protocols previously described by us (Pereira et al., 2017). The genotypes of OPA1 deficient and control mice are as described for satellite cell experiments. Standard chow was fed to mice during their 4-week recovery period following tamoxifen injections at 4 weeks of age. A subgroup of OPA1 deficient and control mice were treated with TUDCA (Millipore: Billerica, MA, USA; 250 mg/kg), administered via intraperitoneal injection. vehicle treated mice received equivalent volumes of phosphate-buffered saline. All animals had free access to standard chow and water in a 22 °C environment with a 12h/12h light/dark cycle.

*Atf4*^fl/fl^ mice were generated as previously described (Ebert et al., 2012). We also generated *Opa1* and *Atf4* double floxed mice from which satellite cells were also isolated.

### Primary Cell Culture

Satellite cell isolation was performed as previously described (Pereira et al., 2017). Eight mice of each experimental condition [control (*OPA1^flox/flox^*) and skeletal muscle OPA1-deficient mice (*OPA1^flox/flox^ has-CreER^T2^)*] were used to isolate primary satellite cells. Satellite cells from *OPA1^fl/fl^*, *Atf4^fl/fl^*, and *OPA1^fl/fl^* + *Atf4*^fl/fl^ double floxed mice were plated on BD Matrigel-coated dishes and differentiated into myoblasts in Dulbecco’s modified Eagle medium (DMEM)-F12 containing 20% fetal bovine serum (FBS), 40 ng/ml basic fibroblast growth factor, 1× non-essential amino acids, 0.14 mM β-mercaptoethanol, 1× penicillin/streptomycin, and Fungizone. Myoblasts were maintained with 10 ng/ml basic fibroblast growth factor and differentiated in DMEM-F12 containing 2% FBS and 1× insulin–transferrin–selenium when 90% confluency was reached. Three days after differentiation, myotubes were infected with an adenovirus expressing GFP-Cre to achieve OPA1 deletion, ATF4 deletion, or double knockout of OPA1 and ATF4. Additionally, we obtained an ATF4 adenovirus (Ad5CMVATF4/RSVeGFP) from the University of Iowa Viral Vector Core facility to induce ATF4 overexpression in Atf4^fl/fl^ myotubes and in OPA1 ablated myotubes.. Experiments were performed between 3 and 7 days after infection.

### Human Cell Culture

Human and primary mouse myotubes were grown and maintained in DMEM-F12 (Gibco: Waltham, MA, USA) supplemented with 20% fetal bovine serum (Gibco), 10 ng/ml basic fibroblast growth factor, 1% penicillin/streptomycin, 300 µl/100 ml Fungizone, 1% non-essential amino acids, and 1 mM β-mercaptoethanol. Before plating, dishes were coated with Matrigel to enhance cell adherence. On alternate days, the cells were washed with phosphate-buffered saline, and the media replaced.

### RNA Sample Preparation and Quantitative RT-PCR in Mouse Samples

RNA was isolated from the gastrocnemius of 40-week-old mice using TRIzol and tissue grinder (Sigma-Aldrich: St. Louis, MO, USA). Reverse polymerase chain reaction (PCR) was performed to generate cDNA using SuperScript III (Invitrogen). For quantitative reverse transcriptase-PCR, 50 ng of cDNA was used for each reaction with iTAQ Universal SYBR green (Biorad: Hercules, CA, USA), using the QuantiStudio 6 Flex system (Applied Biosystems: Waltham, MA, USA). Gene expression was analyzed using the ΔΔC_T_ method, and relative expression was normalized to *Gapdh*. Sequences for primers are listed in supplementary materials (See Table 1).

### RNA Extraction and Quantitative RT-PCR for Cells

RNA extraction and quantification were performed as previously described (Pereira et al., 2017). Briefly, RNA was extracted from samples using TRIzol reagent and purified with the RNeasy kit (Qiagen Inc: Hilden, Germany). The RNA was reverse transcribed, followed by quantitative PCR (qPCR) reactions using SYBR Green. Samples were then loaded onto a 384-well plate, and a qPCR was performed using an ABI Prism 7900HT instrument. The primers used are listed in Supplemental Table 1.

### Western Blotting

To obtain total protein extracts from differentiated myotubes, cells were washed with ice-cold PBS before adding cold lysis buffer (25 mM Tris HCl, pHLJ7.9, 5LJmM MgCl_2_, 10% glycerol, 100LJmM KCl, 1% NP40, 0.3 mM dithiothreitol, 5 mM sodium pyrophosphate, 1LJmM sodium orthovanadate, 50 mM sodium fluoride, and protease inhibitor cocktail (Roche Applied Science: Penzberg, Germany). Cells were scraped, homogenized with a 25-gauge needle, and centrifuged at 14,000 rpm for 10 min at 4 °C. Supernatants were collected and mixed with Laemmli sample buffer to get a final concentration of 1X. Cell lysates were subjected to sodium dodecyl sulfate-polyacrylamide gel electrophoresis (SDS-PAGE), and proteins were transferred to nitrocellulose membranes (BioRad: Berkeley, California, USA). Membranes were blocked with 5% bovine serum albumin (BSA)-Tris-buffered saline with Tween-20 (TBST) and incubated with antibodies as indicated. Quantification was performed using Image Studio Lite Ver 5.2 with three biological replicates for each protein of interest.

Primary antibodies used for Western blotting and their working dilutions include: OPA1 (1:1,000; BD Biosciences: San Jose, CA, USA; #612606), glyceraldehyde 3-phosphate dehydrogenase (GAPDH; 1:5,000; Cell Signaling Technology: Danvers, MA, USA; #2118), MFN1 (1:1,000; Abcam, Cambridge, UK; #ab57602), MFN2 (1:1,000; Abcam; #ab101055), GRP75 (1:1000; Cell Signaling Technology; #D13H4), PACS2 (1:1,000; Abcam; #ab129402), BIP (1:5,000; BD Biosciences; #610978), ATF4 (1:1000; Proteintech: Rosemont, IL, USA; 10835-1-AP), VDAC (1:1000; Cell Signaling Technology; #4866), SERCA (1:1000; Cell Signaling Technology; #4388), MCU (1:1000; ThermoFisher Scientific: Waltham, MA, USA; #PA5-68153), Alpha Tubulin (1:1000; Cell Signaling Technology; #2144), and IP_3_R3 (1:1000; BD Biological Sciences; #610312). Secondary antibodies include: IRDye 800CW anti-mouse (1:10,000; LI-COR: Lincoln, NE, USA; #925-32212) and Alexa Fluor anti-rabbit 680 (1:10,000; Invitrogen; #A27042). Fluorescence was quantified using the LiCor Odyssey imager.

### MERC Confocal Analysis

Cells were grown in culture media on a #1.5 cover glass for optimized microscope optics and either embedded into a petri dish or divided with plastic-walled growth chambers. Cells were incubated for 30 minutes with MitoTracker Orange FM (Invitrogen, 400 nmol/L) and subsequently fixed in 4% paraformaldehyde to prepare for colocalization staining with GRP78. Confocal image stacks were captured with a Zeiss LSM-5, Pascal 5 Axiovert 200 microscope using LSM 5 version 3.2 image capture and analysis software and a Plan-APOCHROMAT 40x/1.4 Oil DIC objective. Images were deconvoluted with NIH (National Institutes of Health) ImageJ software and BITPLANE Imaris software (Oxford Instruments: Abingdon, United Kingdom). Subsequently, images were subjected to analysis using Imaris software for Mander’s and Pearson’s coefficient analyses. Each experiment was performed at least three times, and quantified from 10–20 cells per condition.

### Sample Preparation for Transmission Electron Microscope (TEM) Analysis

Mice were anesthetized with isoflurane according to prior protocols (Garza-Lopez et al., 2022; Lam et al., 2021; Neikirk, Vue, et al., 2023; Vue et al., 2022). Soleus muscle was freshly excised and immediately washed in ice-cold saline. Samples were processed as previously described by us (Pereira et al., 2017). Soleus samples were fixed with Trump’s TEM fixative as previously described(Hinton et al., 2023) and stained. Random images of intermyofibrillar (IMF) sections were obtained. TEM was used to view the ultrastructure of the samples to quantify MERCs.

Cells were fixed in 2.5% glutaraldehyde in sodium cacodylate buffer for 1 hour at 37 °C, embedded in 2% agarose, postfixed in buffered 1% osmium tetroxide, stained in 2% uranyl acetate, and dehydrated with an ethanol-graded series. After embedding in EMbed-812 resin, 80 nm sections were cut on an ultramicrotome and stained with 2% uranyl acetate and lead citrate. Images were acquired on a JEOL JEM-1230 Transmission Electron Microscope, operating at 120 kV.

### TEM Analysis

Measurements of mitochondrial area, circularity, and length were performed using the Multi-Measure region of interest (ROI) tool in ImageJ (Garza-Lopez et al., 2022; Lam et al., 2021; Neikirk, Vue, et al., 2023; Parra et al., 2013; Vue et al., 2022), in accordance with prior recommendations (Neikirk, Lopez, et al., 2023). To measure cristae morphology, we used three distinct ROIs of all the same magnification in ImageJ to determine the cristae area, circulatory index, number, and cristae score. Cristae volume was determined by the sum of all cristae area divided by total area of mitochondria (Patra et al., 2016). The NIH ImageJ software was used to manually trace and analyze all mitochondria or cristae using the freehand told in the ImageJ application (Parra et al., 2013). We also measured MERC distance, length, and percent coverage. MERC length was defined as the horizontal distance between the mitochondria and ER interface. MERC distance was defined as the vertical distance between the mitochondria and ER interface. MERC percentage coverage was calculated as the total ER or mitochondrion surface area normalized to the distance between the mitochondria and the ER.

### Serial Block Face-Scanning Electron Microscopy (SBF-SEM) Processing for *Drosophila* Muscle Fibers

Protocols follow previously established methods (Crabtree et al., 2023; Garza-Lopez et al., 2022; Vue et al., 2022, 2023). Briefly, tissue was fixed in 2% glutaraldehyde in 0.1 M cacodylate buffer and processed using a heavy metal protocol adapted from a previously published protocol (Courson et al., 2021; Mustafi et al., 2014). Tissue was immersed in 3% potassium ferrocyanide and 2% osmium tetroxide (1 hour at 4 °C), followed by filtered 0.1% thiocarbohydrazide (20 min), 2% osmium tetroxide (30 min), and left overnight in 1% uranyl acetate at 4 °C (several de-ionized H_2_O washes were performed between each step). The next day, samples were immersed in a 0.6% lead aspartate solution (30 min at 60 °C and dehydrated in graded acetone (as described for TEM). The tissues were impregnated with epoxy TAAB 812 hard resin, embedded in fresh resin, and polymerized at 60 °C for 36–48 hours. After polymerization, the block was sectioned for TEM to identify the area of interest, then trimmed to 0.5 mm × 0.5 mm and glued to an aluminum pin. The pin was placed into an FEI/Thermo Scientific Volumescope 2 scanning electron microscope (SEM), a state-of-the-art SBF imaging system. For 3D EM reconstruction, 300 to 400 thin (0.09 µm) serial sections were obtained from single blocks that were processed for conventional TEM. Serial sections were collected onto formvar coated slot grids (Pella, Redding CA), stained, and imaged, as described above. The segmentation of SBF-SEM reconstructions was performed by manually tracing structural features through sequential slices of the micrograph block. Images were collected from 30–50 serial sections, which were then stacked, aligned, and assembled into videos using Amira (Thermo Fisher Scientific, Waltham, Massachusetts, USA) to quantify and visualize volumetric structures (Garza-Lopez et al., 2022).

### Proximity Ligation Assay in Cells

We used Duolink In Situ Detection Reagents Red (Sigma-Aldrich; DUO92008) to verify the proximity of two antibodies of interest in cell culture. Previously validated antibodies were used. The first antibody was mouse monoclonal antibody against MFN1 (Abcam #ab57602), the second antibody was a rabbit polyclonal antibody against MFN2 (Abcam #ab50843), produced using a synthetic peptide corresponding to human Mitofusin 2, aa 557–576, conjugated to keyhole limpet hemocyanin (KLH). Both antibodies were determined to work at a 1:200 dilution. Three controls were included in these experiments. Each primary antibody was omitted from separate cells to determine the non-specific binding of the primary antibodies and to confirm the optimal antibody titer. One well was devoid of all primary antibodies to detect any non-specific binding of the Duolink^©^ PLA probes.

Cells were fixed in 4% paraformaldehyde in PBS at 37 °C for 10 minutes. After three rinses with PBS, a permeabilization solution containing 0.02% Triton X-100 in PBS was applied for 10 minutes. A ready-to-use blocking solution from the kit was applied for 1 hour at 37 °C. A cocktail containing both antibodies, or control solution, diluted in the provided diluent, was applied to the cells overnight at 4 °C. The next day, cells were washed and incubated for 1 hour at 37 °C with a pair of oligonucleotide-labeled secondary antibodies, one against mouse and one against rabbit. Cells were washed again and hybridized with the ligase solution for 30 minutes at 37 °C, connecting the oligos in close proximity to each other and forming a closed circle DNA template. Cells were then washed, rolling circle amplification (RCA) was performed with the template, with the probe acting as a primer for the DNA polymerase. RCA was allowed to proceed in a dark hybridization chamber at 37 °C for 1 hour and 40 minutes. Then, cells were washed and mounted in Duolink mounting media and stored at 4 °C overnight and imaged with an SP-8 confocal inverted microscope using a white laser light set to an excitation wavelength of 594 nm and an emission wavelength of 624 nm ± 10 nm.

### Proximity Ligation Assay in Tissue

Duolink In Situ Detection Reagents Red (Sigma-Aldrich; DUO92008) was used to verify the proximity of two antibody locations in tissue samples. We used the two antibodies described for cells. Both antibodies were determined to work at a 1:50 dilution. One tissue slide was devoid of both primary antibodies to detect any non-specific binding of the Duolink PLA probes. The tissues were fixed in a manner that described forlike PLA in cells above except, tissue fixation occurred in warm 4% paraformaldehyde in PBS at 37 °C for 24 hoursinstead of. After fixation, tissues were mounted in optimal cutting temperature (OCT) medium.

### Fly Strains and Genetics

Genetic crosses in *Drosophila melanogaster* were performed on yeast corn medium at 22 °C unless otherwise noted. The *W^1118^* strain was used as a control for the respective genetic backgrounds. *Mef2-Gal4* (III) was used to drive muscle-specific *Opa1-like* (Opa1)(Rai et al., 2014; Vue et al., 2022). *Tub-Gal80^ts^*; *Mef2 Gal4* (BS# 27390) was used for the conditional *Opa1-like* muscle-specific gene KD. Genetic crosses were set up at 18 °C and then shifted to 29 °C at the larval stage. *UAS-mito-GFP* (II chromosome) was used to visualizemitochondria. Stocks were requested or obtained from the Bloomington *Drosophila* stock center. All chromosomes and gene symbols are as described in Flybase (http://flybase.org).

### Mitochondrial Staining

Indirect flight muscles (IFMs) from 2–3-day-old adult flies were dissected and fixed in 4% paraformaldehyde for 1.5–2 hours on a rotor with a minimum rpm. Samples were washed in PBSTx (PBS +0.03% TritonX-100) three times each for 15 min and incubated in PBS with 500– 1000 nM Mitotracker (M22425, Molecular Probes: Eugene, OR, USA; 1 mM stock in dimethyl sulfoxide, DMSO) at 37 °C for 30 min. *DMef2-Gal4* driven *UAS-mito-GFP* was also used to image mitochondria. Muscle samples were washed and counterstained with Phalloidin-TRITC or FITC (P1951-TRITC, Sigma-Aldrich, 50 µg/ml stock; 46950, Sigma-Aldrich) and mounted using Prolong Glass Antifade Mountant with NucBlue (P36985, ThermoFisher, USA). Samples were imaged using a ZEISS 780 confocal microscope and processed using ZEN software (version 3.2.0.115).

### Immunohistochemistry

Two-day-old adult fly thoraces were dissected to obtain IFMs in 4% paraformaldehyde (Sigma), with muscle fixed in 4% paraformaldehyde (Sigma-Aldrich) for 1.5–2 hours at 25 °C on a rotor. Samples were washed three times with PBSTx (Phosphate Saline Buffer with Tween) for 15 min and blocked for 2 hours at 25 °C or overnight at 4 °C using 2% BSA (Sigma-Aldrich). Samples were incubated with respective primary antibodies at 4 °C overnight. Later, samples were washed three times for 10 min with PBSTx and incubated for 2.5 hours in respective secondary antibodies at 25 °C or overnight at 4 °C. Samples were then incubated for 40 min with Phalloidin-TRITC (2.5 µg/ml; P1951, Sigma-Aldrich, USA) to counterstain samples. After incubation for 20 min at 25 °C in Prolong Glass Antifade Mountant with NucBlue stain (P36985, ThermoFisher, USA), samples were mounted. Images were acquired using the ZEISS 780 confocal microscope and processed using ZEN software (version 3.2.0.115). Antibodies used for the staining were rabbit anti-PAC (1:20), rabbit anti-VDAC1 (1:200), rabbit anti-IP3 (1:200), and Alexa Fluor-488-labeled anti-rabbit IgG (1:500, ThermoFisher, USA).

### Electron Microscopy (EM) Sample Preparation in Flies

Flies (2–3 days old) were dissected in standard fixative (2.5% glutaraldehyde, 1% paraformaldehyde, and 0.12 M sodium cacodylate buffer). Individual *Drosophila* muscle fibers (IFMs) were isolated and transferred to a fresh fixative tube. Fixation and imaging were performed as described above.

### RNA Sample Preparation in Flies

RNA was isolated from the IFMs of 1–2-day-old flies. IFMs were removed from the bisected thoraces at 4 °C and immersed in TRIzol (Sigma-Aldrich). Total RNA was extracted in TRIzol reagent (Invitrogen) using a tissue grinder. Reverse polymerase chain reaction was performed to generate cDNA using SuperScript III (Invitrogen). For quantitative reverse transcriptase-PCR, 50 ng of cDNA were used for each reaction with iTAQ Universal Sybergreen (Biorad), using the QuantiStudio 6 Flex system (Applied Biosystems). Gene expression was analyzed using the ΔΔC_T_ method, and relative expression was normalized to GAPDH. Sequences for the primers are listed in Supplementary Table 2.

### ChIP-seq

ChIP-Seq data was analyzed from data sets generated previously (J. Han et al., 2013; Tameire et al., 2019; Zou et al., 2022).

### Statistical Analysis

Results are presented as the mean ± standard error of the mean. Data were analyzed using an unpaired T-test. If more than two groups were compared, one-way analysis of variance (ANOVA) was performed, and significance was assessed using Fisher’s protected least significance difference test. GraphPad PRISM and Statplus software package was used for t-tests and ANOVA analyses (SAS Institute: Cary, NC, USA). For all statistical analyses, a significant difference was accepted when P < 0.05.

## Results

### *Opa1-like* KD in *Drosophila* Flight Muscle Alters Mitochondria, Cristae, and MERC Morphology

Mammalian OPA1 plays an essential role in mitochondrial fusion, and *OPA1* KD increases the number of fragmented mitochondria and the average number of mitochondria per cell (Cipolat et al., 2004; Frezza et al., 2006; Parra et al., 2013; Patten et al., 2014; Scorrano et al., 2002; Tezze et al., 2017; Varanita et al., 2015). We previously reported that *Opa1* downregulation in skeletal muscle of mice induces the mitochondrial fusion and MERC protein MFN2, suggesting that *Opa1* deletion may play a role in regulating MERCs (Pereira et al., 2017). To determine the role of OPA1 in MERC formation, maintenance, and function, we analyzed the effects of *Opa1-like* KD on MERC gene mRNA transcripts in fly muscle tissue (Fig. 1). Using quantitative polymerase chain reaction (qPCR), we find the expression of genes encoding ER and mitochondrial proteins, such as the mitochondrial fission protein dynamin-related protein 1 (*DRP1*) and the ER stress-related markers *ATF4*, *BIP,* and inositol-requiring enzyme 1 (*IRE1*) were increased in *Opa1-like* KD compared with wild-type (WT) *Drosophila* muscles (Fig. 1A) despite the reduced size and activity of the *Opa1-like* KD flies (Supplementary Figure [SF] 1A-B). However, no significant difference in the ER stress marker *ATF6* expression was observed between WT and *Opa1-like* KD muscles (Fig. 1A). Transcripts encoding Ca^2+^-related MERC components, IP_3_R3 (*Itpr3*) and GRP75 (*Hspa9*), were elevated in *Opa1-like* KD fly muscle (Fig. 1A). Interestingly, the gene expression levels of the OMM protein VDAC (Fig. 1A) were decreased. These results indicate that *Opa1-like* KD induces the transcription of MERC components, and ER stress genes.

**Figure 1.**
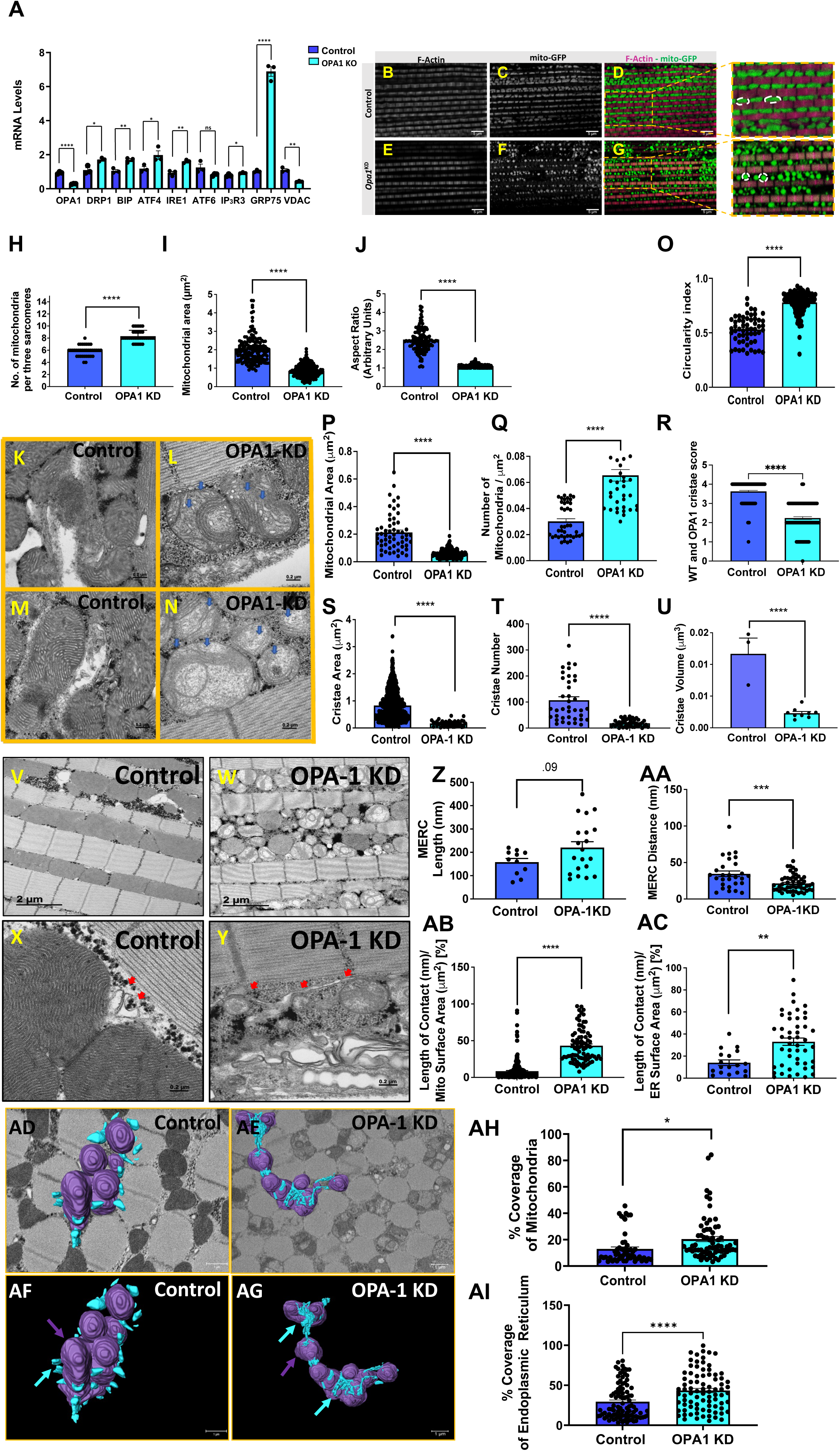
*Opa1-Like* knockdown (KD) in skeletal muscle alters mitochondrial morphology and mitochondrial-ER contact sites in *Drosophila*. (A) mRNA expression levels of genes encoding mitochondria–endoplasmic reticulum (ER) contact (MERC) proteins, *IP_3_R3*, *GRP75* and *VDAC* and ER stress induced proteins in *Opa1-like* KD compared with WT (Control) IFMs. (B-G) Adult myofibrils were stained with Phalloidin-FITC (F-actin) and mito-GFP (mitochondria). IFMs from WT flies revealed mitochondria with tubular morphology, whereas the *Opa1-like* KD flies showed clusters of spherical mitochondria. (H) Mitochondria number (per three sarcomeres) in *Opa1-like* KD (n = 34) and WT (n = 32) IFMs. (I) Quantification of mitochondrial area. (J) Mitochondrial aspect ratio (major axis/minor axis) in *Opa1-like KD* (n = 141) and WT (n = 105) IFMs. (K-N) Transmission electron microscopy (TEM) images showing cristae and mitochondrial morphology in WT and *Opa1-like* KD IFMs (n = 20). (O) Circularity index in *Opa1-like* KD IFMs relative to WT IFMs. (P). Quantification of mitochondrial area in *Opa1-like* KD and WT. (Q) Mitochondria number in *Opa1-like* KD and WT. (R) Cristae quality and abundance was assessed using the cristae score, on a scale from 0 (no sharply defined cristae) to 4 (many regular cristae). The average cristae score was significantly lower in *Opa1-like* KD than in WT. (S) Quantification of cristae area in *Opa1-like* KD and WT. (T) Cristae number in *Opa1-like* KO and WT. (U) Cristae volume in *Opa1-like* KD and WT. (V-Y) TEM images displaying MERCS in WT and *Opa1-like* KD IFMs (n = 20). (Z) Quantification of MERC lengths between *Opa1-like* KD and WT IFMs. (AA) Quantification of MERC distance in *Opa1-like* KD and WT IFMs. (AB) Percentage of mitochondrial surface area in direct contact with the ER in *Opa1-like* KD and WT IFMs. (AC) Percentage of ER surface area in direct contact with mitochondria in *Opa1-like* KD and WT IFMs. (AD-AG) The 3D distribution of single continuous and stationary mitochondria (purple, purple arrows) and ER (blue, blue arrows), reconstructed from serial block face-scanning electron microscopy (SBF-SEM) image stacks of Drosophila indirect flight muscle (IFM) fibers. (AH, AI) Percentage of ER surface area in contact with mitochondria (AH) and percentage mitochondrial surface area in direct contact with ER (AI) in *Opa1-like* KD compared with wild-type (WT) *Drosophila* skeletal muscle (SV1 [WT] SV2-3 [*Opa1-like* KD]). SBF-SEM reconstructions from 7 to 23 fully constructed mitochondria, ER, or MERCs. *Significance was determined by two-tailed Student’s t-test. * P<0.05, ** P<0.01, *** P<0.001, **** P < 0.0001*.

To examine the effects of *Opa1-like* KD on mitochondrial morphology in *Drosophila* skeletal muscle, we used confocal microscopy and transmission electron microscopy (TEM). Confocal microscopy revealed that *Opa1-like* KD increased mitochondrial fragmentation (Fig. 1B-G), as assessed by mitochondrial number normalized to the distance of three sarcomeres (Fig. 1H). Mitochondrial area and aspect ratio (length:width ratio) were also reduced in the *Opa1-like* KD muscle (Fig. 1I-J). Similar results were obtained by TEM (Fig. 1K-N), showing that *Opa1-like* KD increased the mean circularity index, decreased the mean mitochondrial area, and increased the number of mitochondria (Fig. 1O-Q). Additionally, we assessed the abundance and quality of cristae using the cristae score, cristae area, cristae number, and cristae volume, which were all decreased in *Opa1-like* KD mitochondria relative to control mitochondria (Fig. 1R–U). In addition, *Opa1-like* KD was associated with loss of lamellar cristae and increased tubular cristae. The increased mitochondrial fragmentation and alterations in cristae shape are consistent with the many previous reports that OPA1 plays a critical role in remodeling mitochondrial and cristae morphology (Cipolat et al., 2004; Frezza et al., 2006; Parra et al., 2013; Patten et al., 2014; Scorrano et al., 2002; Tezze et al., 2017; Varanita et al., 2015).

After confirming the expected *Opa1-like* KD phenotype [Supplemental Video (SV)1; SV2-3; Figure 1)., TEM was then used to visualize and quantify MERC morphology in *Opa1-like* KD *Drosophila* muscle (Fig. 1V-Y). Relative to WT, *Opa1-like KD* trended toward an increased MERC contact length (Fig. 1Z), decreased the mean MERC distance (Fig. 1AA), increased the percentage of mitochondrial coverage by ER (Fig. 1AB), and increased the percentage of ER coverage by mitochondria (Fig. 1AC), all suggesting that *Opa1-like* deficiency increased MERC tethering. Because TEM only captures structural MERC changes in two dimensions (2D), serial block face-scanning electron microscopy (SBF-SEM), which offers numerous advantages in volumetric renderings (Marshall et al., 2023), was used to validate MERC changes in a three-dimensional (3D) context. The percentage of mitochondrial surface area covered by the ER and the percentage of the ER surface area covered by mitochondria in *Drosophila* muscle both significantly increased in the *Opa1-like* KD relative to the WT muscle (Fig. 1AD-AI, SF 2A-B, SV 4-5). Together, these results demonstrate that OPA1 deficiency increases MERC formation in *Drosophila* skeletal muscle.

### OPA1 Deficiency in Myotubes and Mouse Skeletal Muscle Increases MERC Sites and the Expression of MERC-Tethering Proteins

To examine whether our *Drosophila* findings translated to mammalian muscle cells, we transduced primary myoblast and myotube cell cultures derived from *Opa1* floxed mice with recombinant adenovirus containing the green fluorescent protein gene (Ad-GFP) or Cre recombinase (Ad-CRE) and evaluated MERC protein abundance and morphology. KO of *OPA1* in primary myoblasts (SF 3A) induced ER stress, as indicated by increased BIP and ATF4 protein levels and increased levels of the MERC proteins MFN2 and GRP75 (SF 3B), similar to what was observed in fly muscle. Structurally, increased ER and mitochondrial colocalization in *OPA1* KO myoblasts and myotubes compared to controls was first confirmed by the overlapping fluorescent signals from Mitotracker and GRP78 (SF 3C-S). Additionally, *OPA1* KO cells showed increased MFN2–MFN1 interactions in both primary myoblasts (SV 6-7) and myotubes (SF 3T-Z) as assessed by *in situ* proximity ligation assays (PLA). At the ultrastructural level (SF 3AA-AD, AM-AP; SV 8-9), *OPA1* KO again decreased MERC distance (SF 3AE) and increased the percentage of mitochondria surface covered by ER (SF 3AF, AQ) and the percentage of the ER surface covered by mitochondria in two and three dimensions (2D/3D), respectively (SF 3AG, AR), despite a decrease in mitochondrial size (SF 3AH). Also, similar to results observed in *Drosophila*, OPA1 deficiency in myotubes decreased the number, area, score, and volume of cristae (SF 3AI–AL). These data suggest that OPA1 loss increases MERC protein expression and MERC tethering similarly in mammalian primary myoblasts and myotubes and adult fly muscles.

To determine whether OPA1 regulation of MERCs extends to mature mammalian skeletal muscle, we next investigated the effects of *OPA1* deletion on MERC structure and expression in mice with skeletal muscle-specific inducible knockdown of OPAl (*OPA1* smKO) (Figure 2). Consistent with the fly and cell culture data, OPA1-deficient skeletal muscles from 40-week-old mice, which represent adult mice, showed increased levels of MERC proteins BIP, PACS2, GRP75, MFN2 and MFN1 (Fig. 2A-B). Similarly, mRNA levels of MERC genes as assessed by qPCR were also increased in *OPA1* smKO muscles (Fig. 2C). Additionally, *in situ* PLA (Fig. 2D-I) revealed increased MFN2–MFN1 (Fig. 2F: SV 10-11) and IP_3_R3–VDAC interactions in *OPA1* smKO muscle fibers (Fig. 2I; SV 12–13). TEM analysis (Fig. 2 J-M) showed that MERC distance was decreased (Fig. 2N) and contact between mitochondrial and ER membranes was increased in 2D and 3D images (Fig. 2O-V; SV 14-15) in *OPA1* smKO mice compared to controls. These data demonstrate that MERC formation also increases in mouse skeletal muscle following adult-onset OPA1 deficiency.

**Figure 2.**
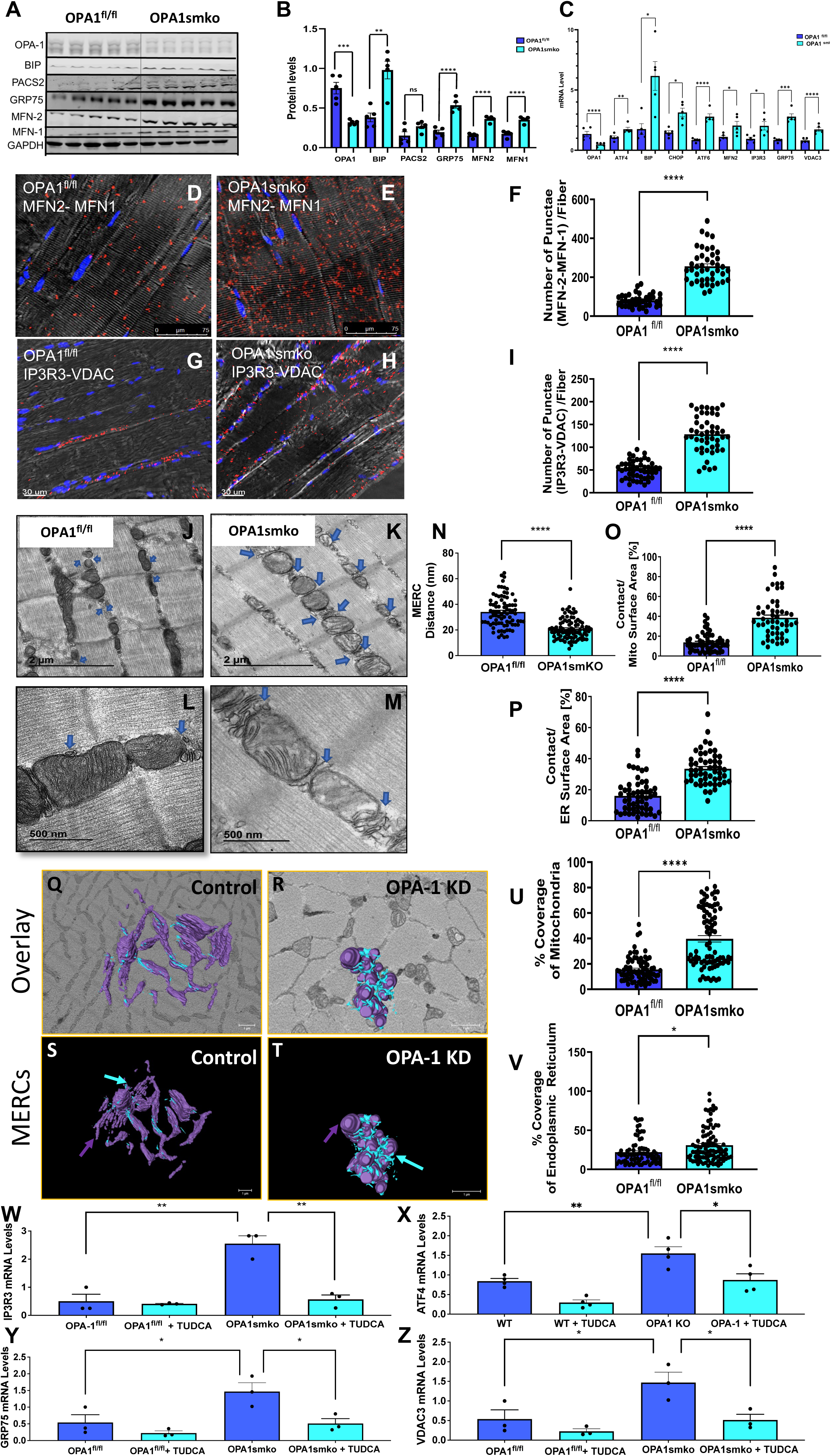
OPA1 deficiency promotes mitochondria–endoplasmic reticulum (ER) contact (MERC) tethering in murine gastrocnemius skeletal muscle. (A) Representative immunoblots showing MERC protein levels (normalized to GAPDH) in 40-week-old WT or *OPA1* smKO muscle (n = 6). (B) Densitometric quantification showing a significant increase in protein levels of BIP, GRP-75, MFN-2, and MFN-1 in *OPA1* smKO compared with WT mice. (C) Quantification of mRNA expression of *OPA1*, ER stress genes *ATF4*, *BIP*, *CHOP*, and *ATF6*, MERC-tethering gene *MFN-2*, and calcium-related MERC genes IP_3_R*3*, *Grp75*, and *VDAC3* in 40-week-old *OPA1* smKO compared with WT mice (n = 5). Data are expressed as fold changes vs. WT mice. (D-I). *In situ* proximal ligation assay (PLA) visualization of MFN1–MNF2 (D-E, red punctae) and IP_3_R3–VDAC interactions (G-H, red punctae) from 40-week-old WT and *Opa1* smKO mice. Quantification demonstrating (F) increased MFN1–MFN2 and (I) IP_3_R3–VDAC interactions in *Opa1* smKO mice compared with WT (n = 3). (J-M). Transmission electron microscopy (TEM) images showing MERCs in 20-week-old wild-type (WT) and *Opa1* skeletal muscle specific knockout (*Opa1* smKO) mice (n = 3). (N). MERC distance in *Opa1* smkO and WT controls. (O, P). Percentage coverage of (O) mitochondrial surface area and (P) ER surface area in *Opa1* smKO relative to WT. (Q-T) The 3D distribution of single continuous and stationary mitochondria (purple, purple arrows) and ER (blue, blue arrows), reconstructed from serial block face-scanning electron microscopy (SBF-SEM) image stacks of *Opa1* skeletal muscle specific knockout (*Opa1* smKO) mouse muscle. (U, V). Percentage of ER surface area in contact with mitochondria (U) and the percentage of mitochondria surface area in direct contact with ER (V) in *OPA1* KO compared with WT skeletal muscle. (W–Z). mRNA levels in 12LJweekLJold *OPA1* smKO and WT mice 4 weeks after TUDCA treatment (n = 3). *IP_3_R3*, (W) *ATF4*, (X) *GRP75 (Y)*, and *VDAC3* (Z) gene expression were significantly reduced in *OPA1*-smKO mice treated with TUDCA compared to those without TUDCA. Data are presented as fold changes vs. WT mice. *SBF-SEM reconstructions from 7 to 23 fully constructed mitochondria, ER, or MERCs. Data are presented as the mean ± SEM. Significance was determined by Student’s t-test. *P < 0.05, **P < 0.01, ***P < 0.001, **** P < 0.0001*.

We previously reported that *OPA1* smKO animals exhibited ER stress activation and increased levels of the MERC protein MFN2 and its interacting partner MFN1 (Pereira et al., 2017). To evaluate the role of ER stress in OPA1-mediated MERC formation, four weeks after tamoxifen injections, *OPA1* smKO mice were treated with either PBS or the chemical ER chaperone tauroursodeoxycholic acid (TUDCA), which prevents unfolded protein response dysfunction (Yoon et al., 2016), for 4 weeks to attenuate ER stress activation (Pereira et al., 2017; Tezze et al., 2017). Following TUDCA treatment, we observed decreased expression of Ca^2+^-related MERC genes *VDAC3*, IP_3_R*3, ATF4*, and *GRP75* in 12-week-old *OPA1* smKO mice relative to untreated *OPA1* smKO mice (Figure 2W–Z). These results support the hypothesis that ER stress may mediate the MERC response to OPA1 deficiency.

### ATF4 Mediates MERC Formation and Spacing in *OPA1*-deficient Skeletal Muscle

To better understand the relationship between ER stress and MERC formation in OPA1-deficient muscles, we hypothesized that the integrated stress response transcription factor ATF4, which is induced in OPA1-deficient muscles and can be activated downstream of the unfolded protein response (UPR), binds to the promoters of MERC genes to induce expression. By performing an *in-silico* study using publicly available ChIP-Seq data, we identified ATF4 binding to the promoters of *Hspa9* (GRP75) and *Vdac3* (SF 4A-B), and potentially two sites in *Itpr3* that encodes IP_3_R3 (SF 4C) in WT samples that are reduced or absent in the ATF4 KO samples. For *Vdac3*, the major peak may be within an intron, suggesting activation downstream of the promoter. These observations support a potential role for stress response factor ATF4 in the observed changes in MERC formation and protein expression under OPA1-deficient conditions.

To further investigate the relationships between ATF4, OPA1, and MERC formation across several models, we evaluated MERC structure and protein abundance in primary mouse myotubes with knockdown or overexpression of ATF4 in concert with OPA1 deficiency (Fig. 3). ATF4 overexpression alone increased the abundance of Ca^2+^-related MERC proteins, IP3R3 and VDAC (Fig. 3A-D), suggesting involvement in MERC formation through ER stress. When we knocked out both OPA1 and ATF4 in muscle cells, the mean mitochondrial area was significantly decreased, whereas the mean mitochondrial area was significantly increased in ATF4 OE muscle cells (Fig. 3E-K) suggesting that ATF4 functions to also modulate mitochondrial fusion. Double OPA1 and ATF4 KO (DKO) and OPA1 KO/ATF4 OE resulted in non-significant reductions in the mitochondrial area relative to control skeletal muscle cells (Fig. 3K), suggesting the pluralistic effects of these proteins serve moderating effects. In contrast to OPA1 KO, ATF4 KO significantly increased MERC distance (Fig. 3L). Conversely, ATF4 OE significantly decreased MERC distance compared to the control (Fig. 3L), indicating that ATF4 regulates the spacing between mitochondrial and ER membranes in myotubes. Moreover, in DKO cells, a significant increase in MERC distance was observed, relative to both control and OPA1 KO cells. These data suggest that OPA1 regulation of MERC spacing is dependent on ATF4 expression.

**Figure 3.**
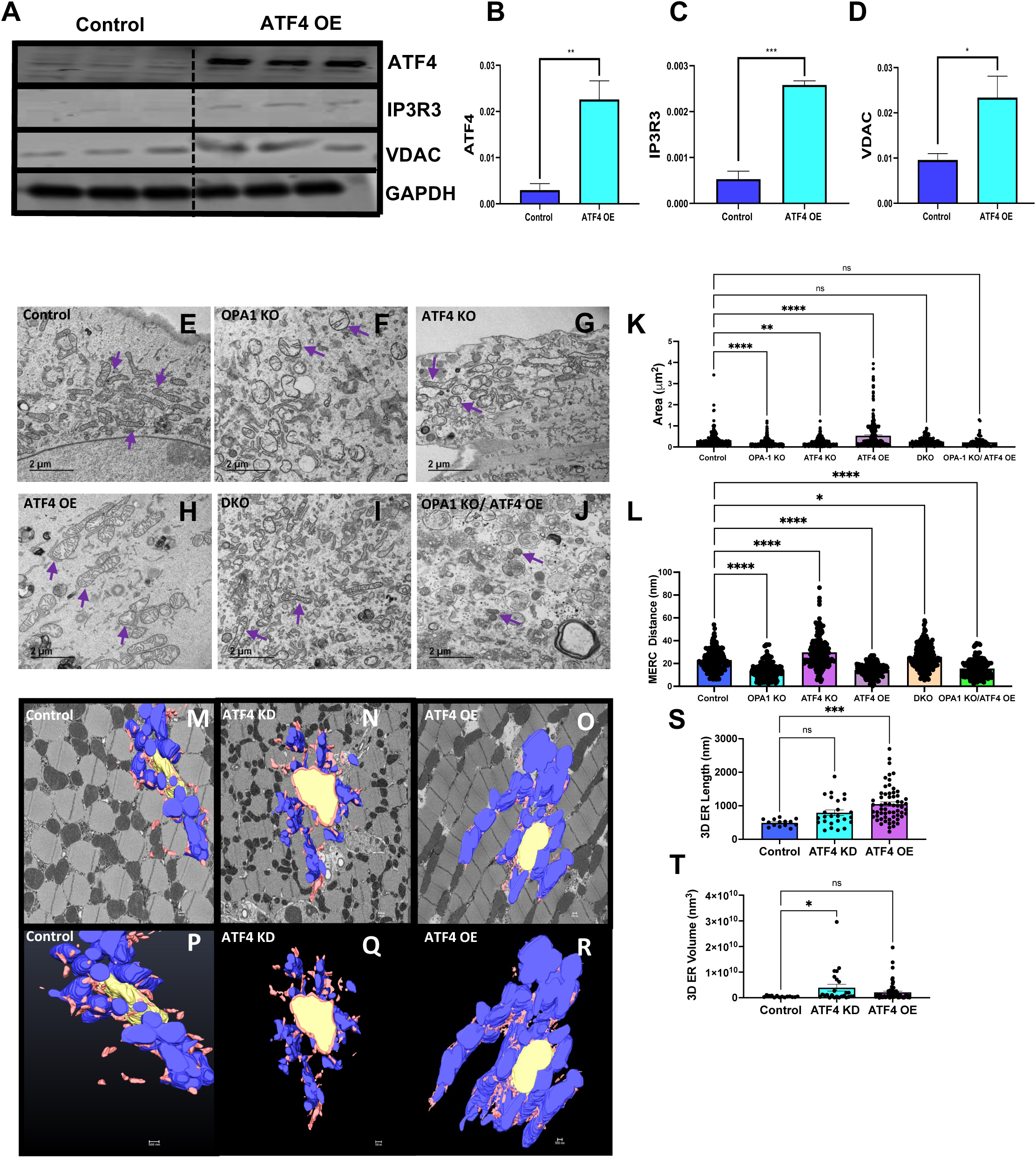
ATF4 regulates mitochondria–endoplasmic reticulum (ER) contact (MERC) formation in *Opa1*-deficient muscle. (A) Representative immunoblot assessing ATF4, IP_3_R3, and VDAC levels from control (with no genetic modifications) or Ad-*Atf4* overexpression (OE) mouse skeletal muscle myotubes (n=6) (B-D). Densitometric quantification demonstrating an increase in (B) ATF4, (C) IP_3_R3 and (D) VDAC protein levels in *Atf4* OE compared with control myotubes. (E-J) TEM panel comparing MERCs area in control, *Opa1* KO, *Atf4* KO, *Atf4* OE, double *Opa1/Atf4* KO (DKO), and *Opa1* KO/*Atf4* OE myotubes. Representative mitochondria are identified with purple arrows. (K) Mitochondrial area in *Opa1* KO, *Atf4* KO, *Atf4* OE, *Opa1/Atf4* KO (DKO), and *Opa1* KO/*Atf4* OE relative to control myotubes. (L) MERC distance in *Opa1* KO, *Atf4* KO, *Atf4* OE, *Opa1/Atf4* KO (DKO), and *Opa1* KO/*Atf4* OE relative to control myotubes. (M-R). Individual electron micrograph of MERCs in control (M), *Atf4* KD (N), and *Atf4* OE myotubes and their serial block face-scanning electron microscopy (SBF-SEM) 3D reconstructions (P-R), showing continuous, and stationary mitochondria (blue), nuclei (yellow), and ER (pink) in *Drosophila* flight muscle. Quantification of ER length (S) and ER volume (T) in control, *Atf4* KD and *Atf4* OE in *Drosophila* flight muscle. *Significance was determined by Student’s t-test. *P < 0.05, **P < 0.01, ***P < 0.001, **** P < 0.0001*.

We next sought to recapitulate these findings in *Drosophila* flight muscle, and SBF-SEM was used to further quantify ER and MERC lengths in *ATF4* KD and *ATF4* OE muscle (Fig. 3M-R, SV 16–18). We found a significant increase in ER length, or length in contact with ER, in *ATF4* OE muscle cells compared with control cells (Fig. 3S), while *ATF4* loss did not alter ER length. The ER volume did not significantly change in *ATF4* OE, but significantly increased in *ATF4* KD cells, likely a byproduct of both increased MERC length and distance (Fig. 3T). Together, these data support the hypothesis that ATF4 expression regulates MERC formation by inducing the expression of MERC proteins, decreasing the width of the MERC space and increasing MERC length and modulating ER morphology. Specifically, a change of both *ATF4* through OE or KD can attenuate OPA1-dependent changes in mitochondrial size, while *ATF4* expression is necessitated for significant MERC tethering.

### OPA1 Deficiency Alters Mitochondrial Ca^2+^ Uptake

Given the alterations in mitochondrial and ER structure observed in relation to OPA1 and ATF4 expression, we sought to explore potential functional consequences of these changes. Prior work suggests that mitochondrial dynamics could play a role in intracellular calcium oscillations in part by regulating MERC formation (CasellasLJDíaz et al., 2021; De Brito & Scorrano, 2008; S. Han et al., 2021). We, therefore, sought to determine changes in Ca^2+^ organellar partitioning in OPA1-deficient myotubes. In cells treated with caffeine (20 mM) to release Ca^2+^ from the ER we estimated mitochondrial Ca^2+^ concentration using mitoPericam fluorescence from control (OPA1^fl/fl^) and OPA1 KO (OPA1^fl/fl^-Cre) primary myotubes (Fig. 4A). The area under the curve was decreased in OPA1 deficient cells, indicating that OPA1deficiency reduces mitochondrial Ca^2+^ uptake (Fig. 4B). The peak amplitude for mitoPericam was also decreased in OPA1 KO, which could implicate reduced mitochondrial calcium uniporter (MCU) activity (Fig. 4C). To further evaluate this possibility, we measured MCU levels in myoblasts, myotubes, and mouse skeletal muscle (SF 5). 4). In all three models, loss of OPA1 increased MCU expression (SF 5B, E, H). To assess ER/SR calcium regulation, levels of the sarcoplasmic/endoplasmic reticulum Ca^2+^-ATPase were measured and noted to be increased upon loss of OPA1 (SF4 C, F, I). Using Fura-2 to assess total cellular calcium, we observed that OPA1 deficiency reduced the area under curve and peak amplitude following caffeine stimulation (Figure 4D-F). When Ca^2+^ re-uptake by the ER was inhibited by thapsigargin (1 µM), both area under the curve and peak amplitude of cytosolic Ca^2+^ were decreased in OPA1-deficient cells following caffeine (20 mM) (Fig. 4G-I). Together these findingsindicate reduced ER Ca^2+^ stores. Thus, OPA1 deficiencyreduces SR calcium, which could contribute to impaired mitochondrial Ca^2+^ uptake, despitecompensatory induction of MCU and SERCA2.

**Figure 4.**
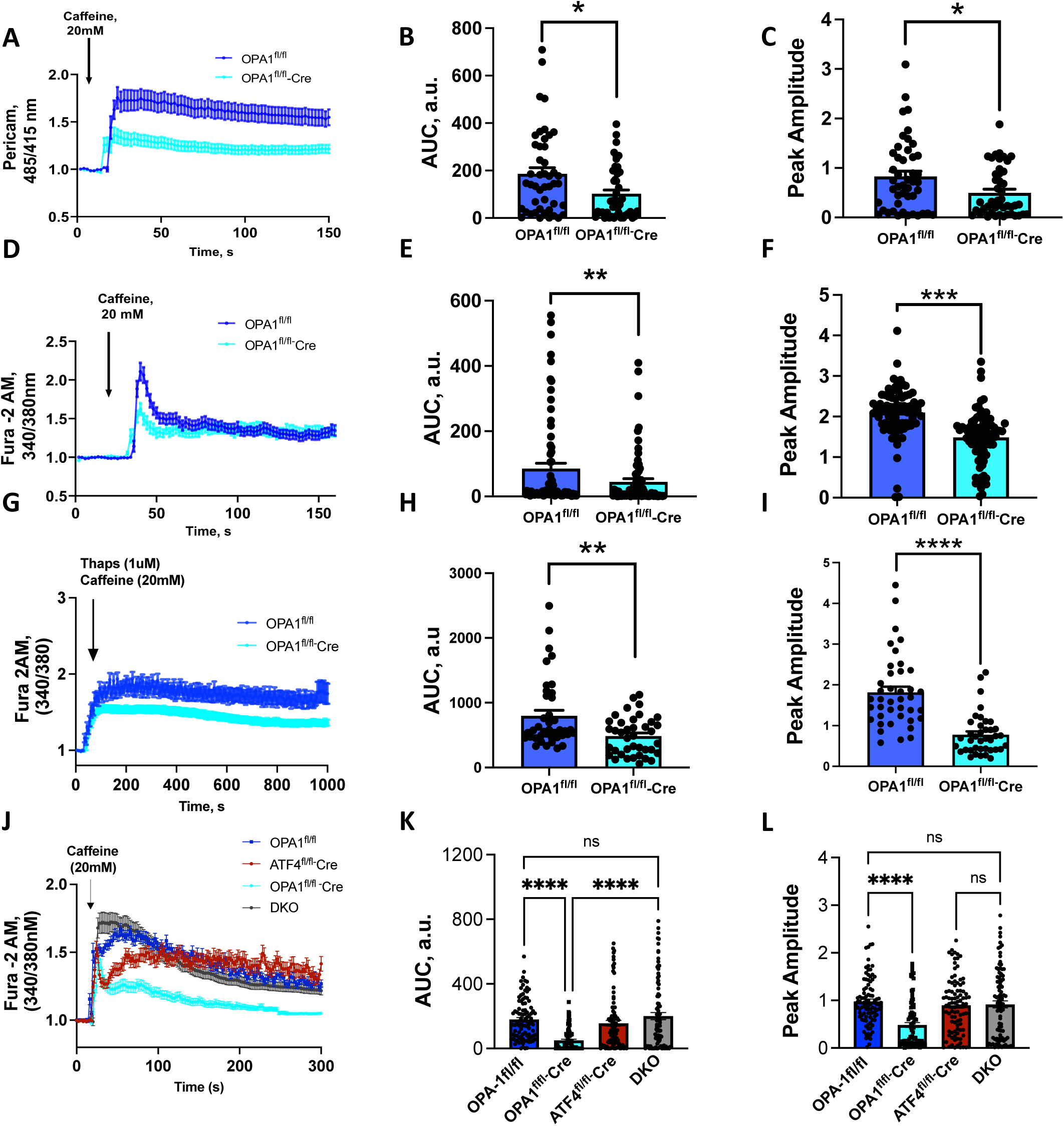
ATF4 regulates calcium homeostasis in OPA1 deficient skeletal myotubes. (A). Mitochondria Ca^2+^ measurements after caffeine administration (20mM) using _mt_pericam in control and *Opa1* KO primary myotubes. (n=3). (B). Area under the curve for pericam showing decreased mitochondrial Ca^2+^ in *Opa1* KO. (C). Quantification of peak amplitude pericam signal showing reduction in OPA1 primary myotubes. (D). Caffeine-induced (20mM) cytosolic Ca^2+^ tracing measured by Fura-2 fluorescence in control and *Opa1* KO primary myotubes(n=5). (E). Decreased area under the curve for cytosolic Ca^2+^in *Opa1* KO myotubes. (F). Quantification of peak Fura-2 amplitude signal in *Opa1* primary myotubes compared to control. (G). Cytosolic Ca^2+^ levels estimated by Fura-2 in control and *Opa1*-KO primary myotubes following treatment with Thapsigargin (Thaps) (1uM) and Caffeine (20mM) (n=5). (H). Area under the curve for Fura-2 fluorescence in *Opa1* KO myotubes compared to control. (I). Quantification of Fura-2 peak amplitude signal in *Opa1* primary myotubes compared to control. (J). Tracing of cytosolic calcium by Fura-2 in primary myotubes from *Opa1* KO, *Atf4* KO, and double KO (DKO) of *Opa1* and *Atf4*. (K). Area under the curve is lower in *Opa1* KO compared control, but normalized in DKO, which is not significantly different relative to control. (L). Peak amplitude quantification significantly decreases in KO of *Opa1* compared to control and is normalized in DKO.

Given the ATF4 dependence of MERC modulation in OPA1 deficient cells described above, we hypothesized that ATF4-dependent MERC formation contributed to altered Ca^2+^ homeostasis and reducing ATF4 could restore Ca^2+^ homeostasis in cells lacking OPA1. We compared cytosolic Ca^2+^ measurements using Fura-2 from control, OPA1 KO and OPA1/ATF4 DKO primary myotubes following caffeine exposure (Fig. 4J). Loss of OPA1 reduced the area under the curve (Fig. 6K) and the peak amplitude, consistent with reduced Ca^2+^ release from the ER into the cytosol (Fig. 4L). However, the loss of ATF4 alone or the simultaneous loss of ATF4 and OPA1 restored area under the curve and peak amplitude to levels similar to those seen in control mice (Fig. 4J-L). This shows that in DKO, ATF4 can rescue impaired calcium homeostasis caused by OPA1 deficiency. These data suggest that ATF4-induced MERCs remodeling contributes to dysregulated Ca^2+^ homeostasis in OPA1-deficient muscle cells (Fig. 4J-L).

## Discussion

Previous studies have used 3D reconstruction to better elucidate diverse phenotypes of mitochondria and ER communication modalities, including the description of curved sheets of rough ER around mitochondria referred to as wrappER (Anastasia et al., 2021). However, to our knowledge, no studies have yet investigated the changes incurred by the loss of OPA1 on MERC formations using a mixed-technique approach including 3D reconstruction. Numerous studies have shown that Protein kinase RNA-like ER Kinase (also known as PERK) is a facilitator of MERC tethering during ER-stress (van Vliet & Agostinis, 2016; Verfaillie et al., 2012). OPA1 function has been linked to the ER in numerous ways, including facilitating calcium transfer and homeostasis (Cartes-Saavedra et al., 2022). Thus, our study supports earlier work demonstrating a link between OPA1 function and the regulation of MERCs. This study advances the field by demonstrating the central role of ATF4 in regulating MERC formation in the context of altered OPA1 expression.

MERCs mediate important intracellular communication processes between the ER and the mitochondria, including Ca^2+^ and lipid transfer that could influence mitochondrial morphology and apoptotic signaling (Giacomello & Pellegrini, 2016; Moltedo et al., 2019; Sassano et al., 2022). MERC defects are associated with the pathophysiology of various disorders including neurodegenerative illnesses, obesity, and diabetes (Kim & Roy, 2020; Liao et al., 2017; Pereira et al., 2017; Tubbs et al., 2018; Verma et al., 2016). Evaluating mechanisms that regulate MERC formation and maintenance has begun to elucidate the role of MERCs in cellular physiology and metabolism. However, the relationship between mitochondrial dynamics, mitochondrial stress, and MERCs remained incompletely understood. Here, we demonstrate that OPA1 deficiency increased key MERC proteins, including MFN2, MFN1, PACS2, and BIP that paralleled bioenergetics defects. OPA1-deficiency also altered MERC tethering at the ultrastructural level and increased MERC volume and percent coverage in skeletal muscle, as determined by SBF-SEM. Importantly, we also elucidated that ATF4 might regulate MERC gene expression in response to mitochondrial stress, which could represent an initial compensatory mechanism that becomes persistent and ultimately maladaptive.

Reduced OPA1 expression results in increased expression of the ER stress protein ATF4, and of MFN2, a tethering MERC protein that regulates Ca^2+^ levels in the MERC space (Ainbinder et al., 2015), (De Brito & Scorrano, 2008; Naon et al., 2016; Pereira et al., 2017; Sood et al., 2014). MERC thickness is thought to be precisely maintained to ensure normal Ca^2+^ transport, as an insufficient distance can result in steric hindrance among various components of the Ca^2+^ transporter machinery (Giacomello & Pellegrini, 2016). Ca^2+^ uptake between the ER and mitochondria is more likely to occur when the organelles are in close proximity, with an ideal distance ranging from 15–30 nm (Csordás et al., 2018; Giacomello & Pellegrini, 2016; Thoudam et al., 2019). Reduced OPA1 expression decreased MERC distance, induced structural changes in MERCs, and were correlated with an increase in the MERC proteins IP3R, GRP75, and VDAC3 and are crucial for Ca^2+^ transfer between the two organelles. Interestingly, despite this reduction in MERCs distance and increased MERC protein levels, we confirmed prior results (Cartes-Saavedra et al., 2020) showing mitochondrial Ca^2+^ uptake was reduced in OPA1-deficient myotubes. The discrepancy between the expression of proteins that promote mitochondrial Ca^2+^ uptake (including MCU), versus the measured reductions in SR and mitochondrial calcium content, underscore additional mechanisms through which OPA1 loss impairs cellular Ca^2+^ homeostasis. Examining decreased store-operated Ca^2+^ entry during OPA1 loss may offer greater insight into Ca^2+^ availability. The induction of the ATF4-dependent pathways identified, could represent an adaptive response. We validated previous results (Cretin et al., 2021; Parra et al., 2013) showing that loss of OPA1 increases mitochondrial fragmentation in concert with the induction of *Drp1,* a regulator of mitochondrial fission (Favaro et al., 2019; Möpert et al., 2009). Mitochondrial size is also known to influence calcium uptake (Golic et al., 2014; Handran et al., 1997; Kowaltowski et al., 2019), and represents an additional mechanism by which OPA1 deficiency could impair intracellular calcium homeostasis.

Changes in MERC properties may also modulate ER to mitochondrial Ca^2+^ transfer, which serves an essential role in the regulation of mitochondrial energy metabolism and oxidative phosphorylation (Bravo et al., 2011; Glancy & Balaban, 2012; Rieusset, 2018). Mitochondrial Ca^2+^ concentration regulates the activation of the TCA cycle, and increased Ca^2+^ levels in the mitochondrial matrix activate multiple enzymes involved in energy production (Carreras-Sureda et al., 2019; Denton et al., 1975; Glancy et al., 2013; Ivannikov & Macleod, 2013; Traaseth et al., 2004). Thus, the reduced intramitochondrial Ca^2+^ levels could contribute to the bioenergetics impairment and altered crista morphology we found to be associated with OPA1 deficiency. Altered MERC morphology in skeletal muscle following OPA1 loss could represent a compensatory mechanism to restore Ca^2+^ homeostasis, given MERCs are understood to serve as regulatory hubs in ER and mitochondrial Ca^2+^ homeostasis (Cartes-Saavedra et al., 2020, 2021; Fülöp et al., 2011; Kushnareva et al., 2013). Similarly, the induction of IP_3_R and SERCA could also represent an adaptive responses. The inhibition of IP_3_R mimics sustained ER stress by impairing mitochondrial function, and inducing autophagy (Jia et al., 2019; Rashid et al., 2015). Moreover, SERCA induction may improve mitochondrial quality control (Tan et al., 2020), offering clues as to the pluralistic roles ATF4 expression has on mitochondrial structure in addition to only MERC formation. However, we cannot exclude the possibility that the associated ATF4-dependent MERCs changes observed in response to OPA1 deficiency could be linked to the impairment of mitochondrial and SR calcium homeostasis via additional mechanisms.

Changes in mitochondria–ER cross-talk accompany the early stages of ER stress, suggesting that exchange in metabolites between the mitochondria and ER are altered in response to various cellular insults (Bravo et al., 2011). The ER-stress-associated protein IRE1α, which was induced in *Opa1-like* KD flies in our experiments, has non-canonical functions in MERC regulation, Ca^2+^ flux between the ER and mitochondria, and in regulating mitochondrial respiration(Carreras-Sureda et al., 2019), indicating a cloe association between ER stress, MERC formation, and the regulation of mitochondrial metabolism.

OPA1 deficiency in murine gastrocnemius skeletal muscle induced ER stress-related proteins, including the unfolded protein response (UPR) transcription factor ATF4, ATF6, and CHOP(Cawley et al., 2011). Recent studies have described that the UPR can also modulate and restore mitochondrial proteostasis which includes reducing unfolded protein load (Hetz, 2012), representing an adaptive response to relieve ER stress (Rainbolt et al., 2014). ATF6-deficiency has been linked to dysfunction in mitochondrial potential, homeostasis, and biogenesis (Wang et al., 2018). Similarly, in neuroblastoma models, IRE1α-XBP1 inhibition decreases MERC spaces (Chu et al., 2021). NRF2, another transcription factor induced by OPA1 deletion, plays pleiotropic roles in mitochondrial biology (Buttari et al., 2022). CHOP which is induced during ER stress might trigger mitochondria-dependent apoptosis (Hu et al., 2019). Thus, many lines of evidence support a link between the UPR transcription factors and mitochondrial function. Our *in-silico* study confirmed that ATF4 binds to downstream factors of MERC genes encoding GRP75 and VDAC3. ATF4 induction following OPA1 deficiency may be an adaptive mechanism for directly regulating MERC gene expression as a means of modifying MERC formation. It is therefore possible that ATF6, XBP1, CHOP, and NRF2 also regulate MERC gene expression by a similar mechanism. Previously the regulation of MERCs by the UPR has been suggested (Amodio et al., 2021; Wilson & Metzakopian, 2021), yet other studies have shown that MERCs can suppress proteins associated with the UPR (Li et al., 2023), underscoring the need for additional studies into this interplay.

In summary, using a model of OPA1-deficiency-induced mitochondrial and ER stress we identified a novel role for ATF4 induction in regulating MERC formation. OPA1 deficiency induces mitochondrial dysfunction that leads to complex compensatory mechanisms, including altered MERC structure and function and perturbed Ca^2+^ homeostasis. Our data suggest that these mechanisms are, at least in part, mediated by ATF4, which regulates ER-mitochondria coupling and calcium homeostasis in skeletal muscle cells likely via transcriptional regulation of core MERC proteins and additional undetermined mechanisms.

## Collaboration Acknowledgements

We thank Dr. Christopher M. Adams for providing the Ad5CMVATF4/RSVeGFP and ATF4 floxed mice. We would like to also thank George R. Marcotte for his assistance in optimizing the Ad5CMVATF4/RSVeGFP for myotubes.

## Funding Acknowledgements

This work was supported by NIH grants R01HL108379 and R01DK092065 to E.D.A; Division of Intramural Research of the National Heart, Lung, and Blood Institute (NHLBI, 1ZIAHL006221 and the National Institute of Arthritis and Musculoskeletal and Skin Diseases (NIAMS) to B.G.; The United Negro College Fund/Bristol-Myers Squibb E.E. Just Faculty Fund, Burroughs Wellcome Fund Career Awards at the Scientific Interface Award, Burroughs Wellcome Fund Ad-hoc Award, National Institutes of Health Small Research Pilot Subaward to 5R25HL106365-12 from the National Institutes of Health PRIDE Program, DK020593, Vanderbilt Diabetes and Research Training Center for DRTC Alzheimer’s Disease Pilot & Feasibility Program to A.H.J.; National Institute of Health (NIH) NIDDK T-32, number DK007563 entitled Multidisciplinary Training in Molecular Endocrinology to Z.V.; NIH NIA RF1AG55549, NIH NINDS R01NS107265, RO1AG062135, AG59093, AG072899 to E.T.; United Negro College Fund/Bristol-Myers Squibb E.E. Just Postgraduate Fellowship in the Life Sciences Fellowship to H.K.B.; and NSF grant MCB #2011577I and NIH T32 5T32GM133353 to S.A.M. Its contents are solely the responsibility of the authors and do not necessarily represent the official view of the NIH. The funders had no role in study design, data collection and analysis, decision to publish, or preparation of the manuscript.

## Supplementary Materials

**Supplemental Figure 1.** Reduced pupal size and reduced mobility in *Opa1-like* knockdown (KD) (A). Representative images of pupa showing reduced size of *Opa1-like* KD compared with WT (control) fly. (B). The number of steps per second were significantly decreased in adult *Opa1-like* KD flies compared with WT. *Data are presented as the mean ± SEM. Significance was determined using Student’s t-test. ****P < 0.0001*.

**Supplemental Figure 2.** Serial block face-scanning electron microscopy (SBF-SEM) demonstrates that the loss of *Opa1* increases mitochondria–endoplasmic reticulum (ER) contacts (MERCs). MERCs are shown in white and mitochondria are purple in wild type (left) and *Opa1* deficient (right) in (A-B) *Drosophila* muscle, (C-D) primary skeletal muscle myotubes, and (E-F) murine muscle.

**Supplemental Figure 3.** OPA1 knockout (*Opa1* KO) increases mitochondria–endoplasmic reticulum (ER) contact (MERC) formation in primary skeletal myoblasts and myotubes. (A) Representative immunoblots of ER stress proteins, ATF4 and BIP, and MERC-tethering proteins, MFN-2 and GRP-75, after infecting *Opa1^fl/fl^* myotubes with Ad-Cre (*Opa1^fl/fl^*-Cre). (B) Densitometric quantification demonstrating an increase in BIP, ATF4, MFN2, and GRP75 levels in *Opa1^fl/fl^*-Cre myotubes compared with WT, normalized to alpha tubulin. (C-J). Representative confocal images of *Opa1^fl/fl^* and *Opa1^fl/fl^*-Cre myoblasts (C-J) and myotubes (K-P), with DAPI-labeled nuclei (blue) (C,G), mitotracker-labeled mitochondria (red) (D,H,K,N), GRP78 (E, I, L, O) showing the ER (green), and a composite image showing all channels (F, J, M, P; n = 10 cells). (Q) Co-localization (Pearson’s correlation coefficient) is increased for ER and mitochondria in *Opa1* KO compared with WT myoblasts (R, S). Mander’s overlap coefficient, demonstrating increased co-occurrence of ER and mitochondria in *Opa1^fl/fl^*-Cre (R) myoblasts (S) and myotubes compared with WT. (T-Y) *In situ* proximity ligation assay (PLA) showing MFN1–MFN2 interactions (red punctae) in *Opa1^fl/fl^*and *Opa1^fl/fl^*-Cre myoblasts (n = 10 cells, 3 independent experiments). Nuclei are labeled with 4′,6-diamidino-2-phenylindole (DAPI, blue). (X) Quantification showing increased MFN1–MFN2 puncta in *Opa1* KO compared with WT myoblasts. (AA-AD) Transmission electron microscopy (TEM) images of MERCs in primary *Opa1^fl/fl^* and *Opa1^fl/fl^*-Cre myotubes (Red arrows indicate mitochondria–ER interactions; n= 3 of 10 cells/experiment). Scale bar = 2 µm (2.5K magnification) and 500 nm (15K magnification). (AE) Quantification of MERC distance in *Opa1^fl/fl^*and *Opa1^fl/fl^*-Cre myotubes. (AF, AG). (AF) Percentage of mitochondrial surface area in direct contact with ER (AG) and percentage of ER surface area in direct contact with mitochondria in *Opa1^fl/fl^*-Cre compared with *Opa1^fl/fl^* myotubes. (AH) MERC area in *Opa1^fl/fl^*-Cre compared with *Opa1^fl/fl^* myoblasts and myotubes. (AI-AL). Cristae number (AI), area (AJ), score (AK) and volume (AL) in *Opa1^fl/fl^*-Cre compared with *Opa1^fl/fl^*. (AM-AP) The 3D distribution of single continuous and stationary mitochondria (purple, purple arrows) and ER (blue, blue arrows), reconstructed from serial block face-scanning electron microscopy (SBF-SEM) image stacks of *Opa1^fl/fl^*-Cre primary myotubes. (AQ-AR). Percentage of ER surface area in contact with mitochondria (AQ) and the percentage of mitochondria surface area in direct contact with ER (AR) in *Opa1^fl/fl^*-Cre compared with *Opa1^fl/fl^* myotubes. SBF-SEM reconstructions from 7 to 23 fully constructed mitochondria, ER, or MERCs. *Data are presented as the mean ± SEM. Significance was determined by Student’s t-test. *P < 0.05, **P < 0.01, ***P < 0.001, **** P < 0.0001*.

**Supplemental Figure 4.** ATF4 binds to the promoters of genes encoding the IP_3_R-GRP75-VDAC MERC Complex. ChIP-seq analysis (represented in blue for the reverse strand and red for the forward strand) from WT and *ATF4* deficient skeletal-muscle derived cells at (A) *Hspa9* (GRP75), (B) *VDAC3*, and (C) *Itpr3* (IP_3_R3) gene loci. Data obtained from a publicly available database: https://chip-atlas.org. Data displays wild-type fibroblast and ATF4 knockout fibroblasts, with ATF4 target genes binding peaks within 3kb from the Transcription Start Site of the annotated gene. The mRNA-Seq data (yellow) reveals the gene expression levels, allowing for gene expression relative to binding to be considered.

**Supplemental Figure 5.** Loss of *Opa1* in myotubes and skeletal muscle increases Mitochondrial Calcium Uniporter (MCU) and Sarcoplasmic/Endoplasmic Reticulum Ca^2+^-ATPase (SERCA) levels. (A). Western blot showing MCU and SERCA levels following OPA1 KO in myoblasts. (B,C) Densitometry for MCU and SERCA respectively, normalized to alpha-tubulin. (D) Western blot showing MCU and SERCA levels following OPA1 KO in myotubes. (E,F) Densitometry for MCU and SERCA respectively. (G) Western blot showing MCU and SERCA levels following OPA1 KO in mouse skeletal gastrocnemius muscle. (H,I) Densitometry for MCU and SERCA respectively.

**Supplemental Table 1.** qPCR primers used for all studies involving mouse samples.

**Supplemental Table 2.** qPCR primers used for all studies involving *Drosophila*.

**Supplemental Videos 1–3.** Loss of *Opa1-like* in *Drosophila* skeletal muscle decreases muscle movement. Live videos of fly movement for WT (Video 1) and *Opa1-like* KD (Video 2 and 3) flies.

**Supplemental Videos 4-5.** Representative 360° rotational view of mitochondria–endoplasmic reticulum (ER) contacts (MERCs) in WT (Video 4) *Drosophila* and after *Opa1-like* knockdown (Video 5), captured with serial block face-scanning electron microscopy (SBF-SEM). Mitochondria are labeled blue, and ER are labeled pink.

**Supplementary Videos 6–7.** Representative 360° rotational view of confocal images in primary myoblasts after *OPA1* knockdown. Nuclei are labeled blue and MFN2–MFN1 interactions are labeled red. Supplemental Video 4 shows OPA1^fl/fl^ while Supplemental Video 5 shows OPA-1^fl/fl^Cre.

**Supplemental Videos 8–9.** Representative 360° rotational view of mitochondria–endoplasmic reticulum (ER) contacts (MERCs) in primary myotubes as WT (Video 8) and after *OPA1* knockdown (Video 9), captured with serial block face-scanning electron microscopy (SBF-SEM). Mitochondria are labeled blue, and ER are labeled pink.

**Supplementary Videos 10-13.** Representative 360° rotational views of confocal images of mouse skeletal muscle. Muscle fibers are labeled gray, and nuclei are labeled blue. Protein antibody interactions, MFN1–MFN2 and IP_3_R3–VDAC interactions, are labeled in red. MFN1– MFN2 interactions are shown in WT (Video 10) and OPA1 smKO (Video 11). IP_3_R3–VDAC interactions are shown in WT (Video 12) and OPA1 smKO (Video 13).

**Supplemental Videos 14-15.** Representative 360° rotational view of mitochondrial–endoplasmic reticulum (ER) contacts (MERCs) in mouse skeletal tissue as WT (Video 14) and after *OPA1* knockdown (Video 15), captured using serial block face-scanning electron microscopy (SBF-SEM). Mitochondria are labeled blue, and ER is labeled pink.

**Supplemental Videos 16–18.** Representative 360° rotational view of mitochondria–endoplasmic reticulum (ER) contacts (MERCs) in *Drosophila* flight muscle from control (Video 16), *ATF4* KO (Video 17), and *ATF4* overexpression (Video 18), captured with serial block face-scanning electron microscopy (SBF-SEM). Mitochondria are labeled blue, ER is labeled pink, and the nucleus is labeled yellow.

