## Supplemental Figures for "ATF4 Dependent Increase in Mitochondrial-Endoplasmic Reticulum Tethering Following OPA1 Deletion in Skeletal Muscle"

### Supplemental Figure 1

A

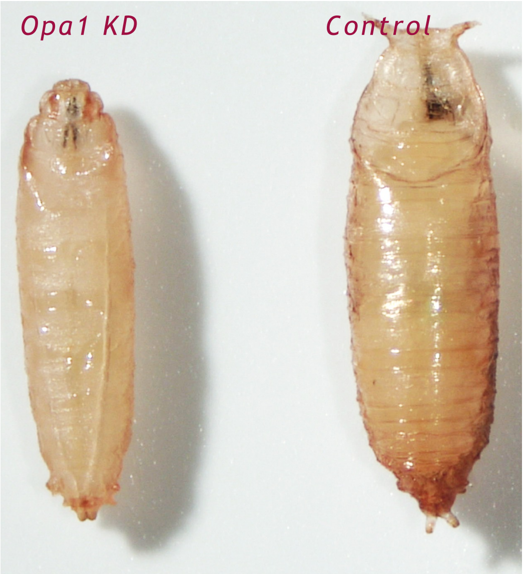

B

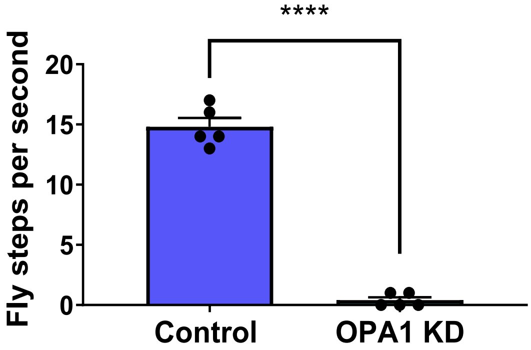

### Supplemental Figure 2

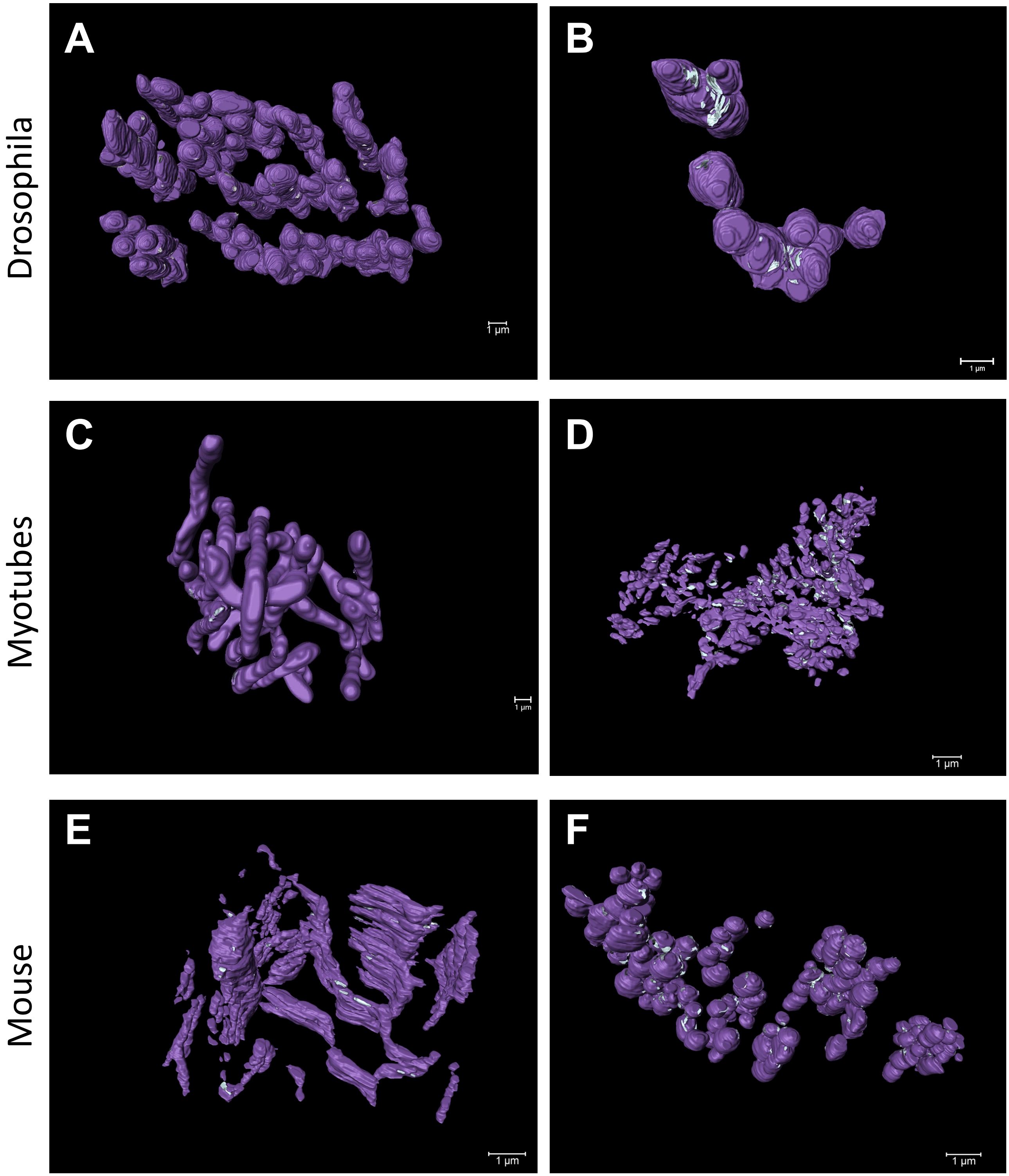

### Supplemental Figure 3

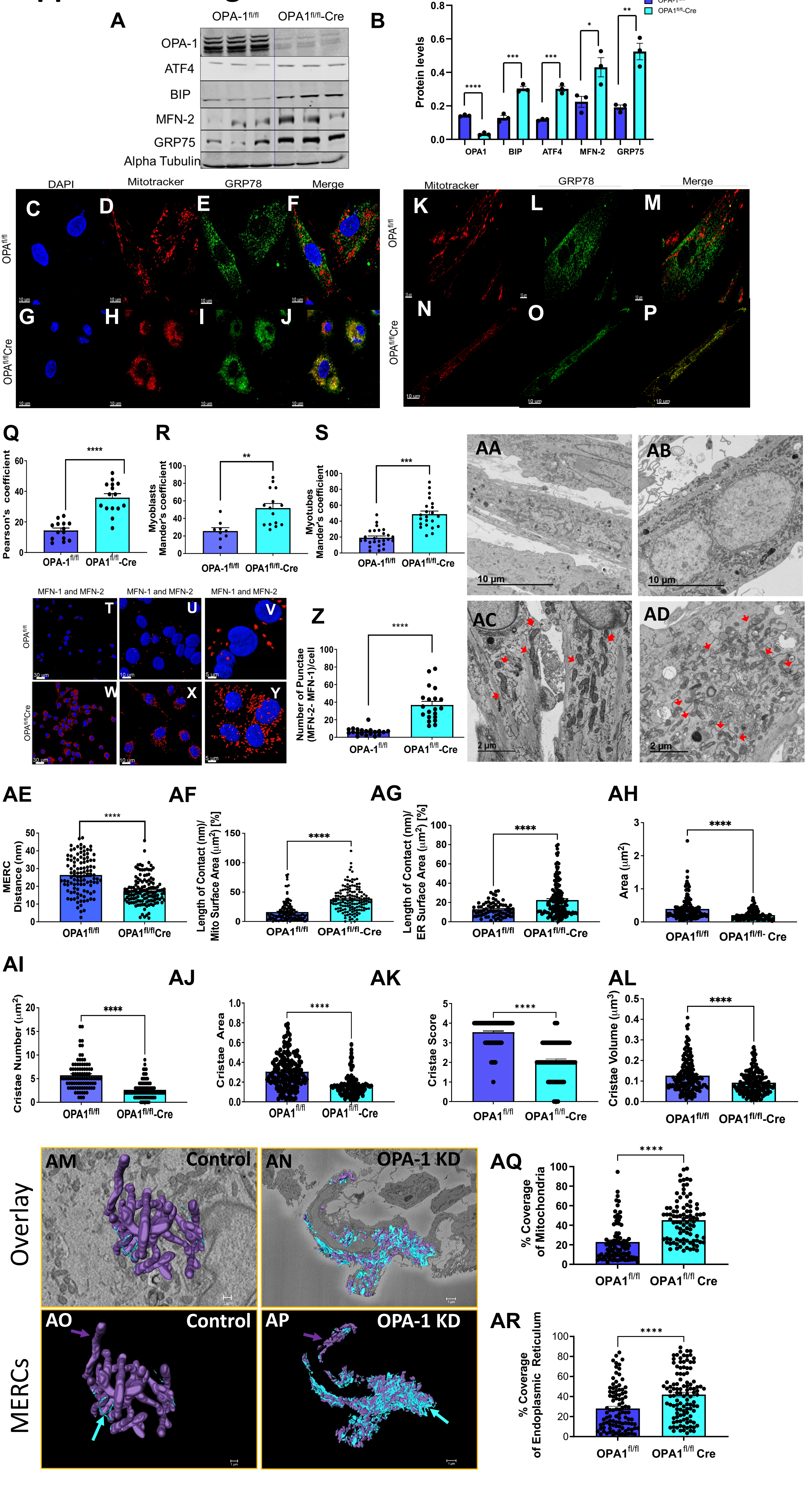

### Supplemental Figure 4

A

GRP75

B

VDAC3

C

$IP_3R$

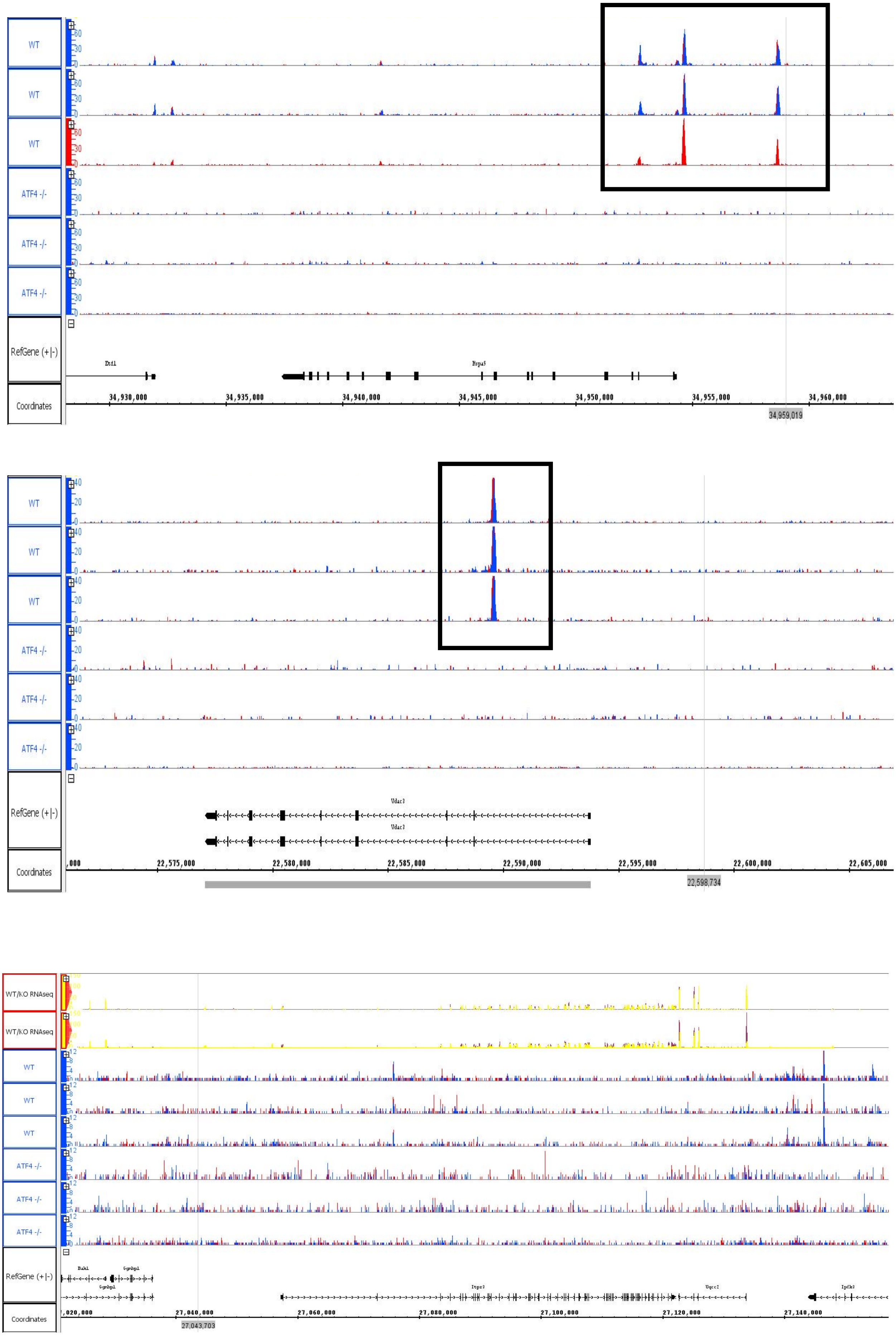

### Supplemental Figure 5

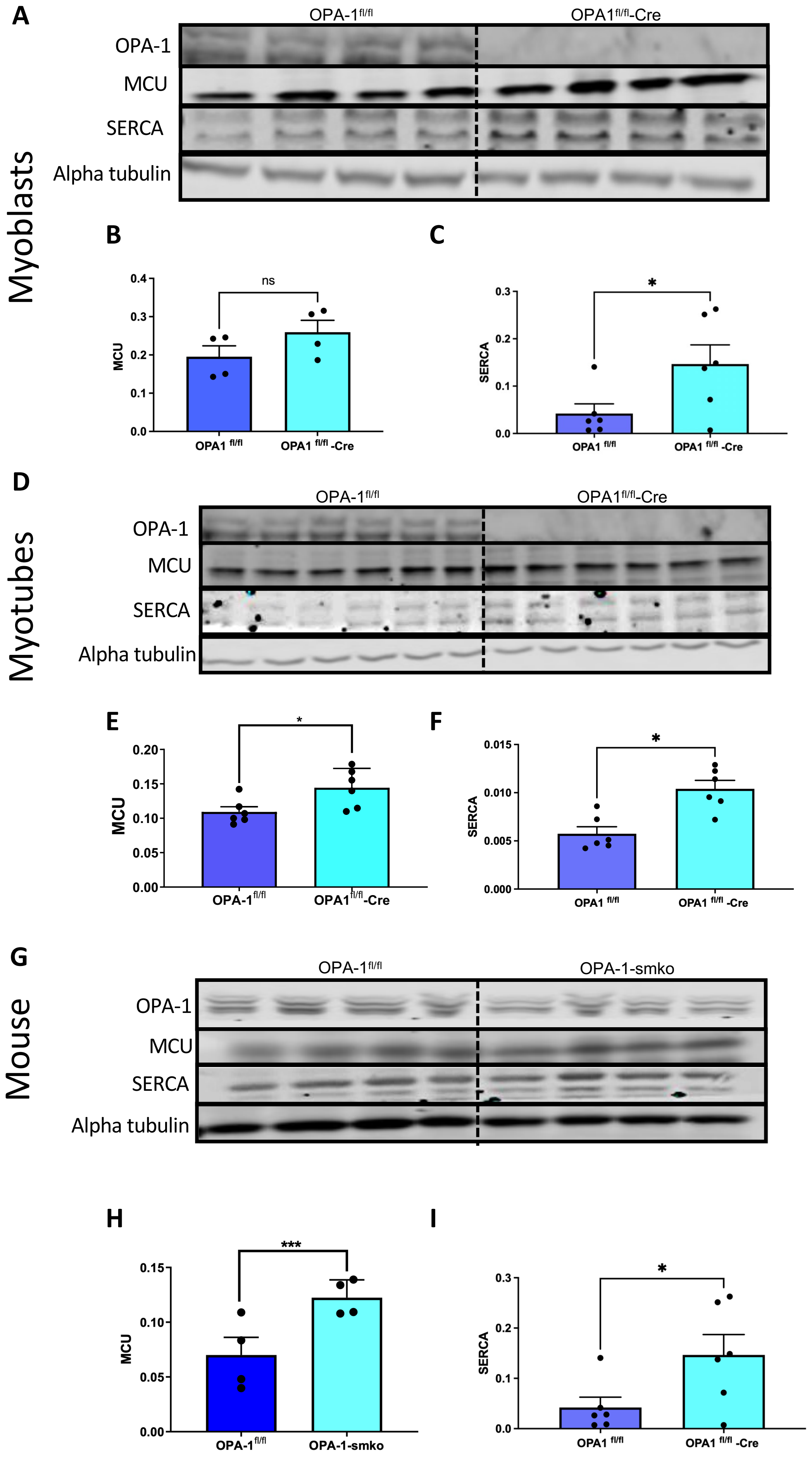

### Table 1. qPCR Primers for Mouse Study

| Gene | Primers | |
|---|---|---|
| CHOP | Forward | 5'GTCCCTAGCTTGGCTGACAGA-3' |
| | Reverse | 5' TGGAGAGCGAGGGCTTT-3' |
| OPA1 | Forward | 5' ACCAGGAGACTGTGTCAA-3' |
| | Reverse | 5'TCTTCAAATAAACGCAGAGGTG-3' |
| ATF4 | Forward | 5'-AGCAAAACAAGACAGCAGCC-3' |
| | Reverse | 5'-ACTCTCTTCTTCCCCCTTGC-3' |
| BiP | Forward | 5'-TCATCGGACGCACTTGGAA-3' |
| | Reverse | 5'-CAACCACCTTGAATGGCAAGA-3' |
| Mouse Gapdh | Forward | 5'ACTCTCTTCTTCCCCCTTGC |
| | Reverse | 5'TCCACGACATACTCAGCAC-3' |
| MFN2 | Forward | 5' gccggacgagcaatgggaag-3' |
| | Reverse | 5'-atggagggcatgggccagtt-3' |
| GRP75 | Forward | 5'-ATGGCTGGAATGGCCTTAGC |
| | Reverse | 5'-GCACCCTTGATTGCTTCTGAT-3' |

### Table 2. qPCR Primers for Drosophila Study

| Gene | Primers | |
|---|---|---|
| RP49 | Forward | 5′-AGGCCCAAGATCGTGAAGAA-3′ |
| | Reverse | 5′-TCGATACCCTTGGGCTTGC-3′ |
| Drp1 | Forward | 5′-TCCACAATCTTCTCGTGCAG-3′ |
| | Reverse | 5′-CATTCACGAGGAGATGCAGC-3′ |
| Marf | Forward | 5′-GTATGTCCATGAGACGACCA-3′ |
| | Reverse | 5′-CTTGTACACATAGCTTTCGA-3′ |
| Opa1 | Forward | 5′-GACTCTGACCGGGAATACGA-3′ |
| | Reverse | 5′-CTAGAACCATGCGTTGCAGA-3′ |
| ATF4 | Forward | 5′-TCGCAAAAGTTGGTTAAACG-3′ |
| | Reverse | 5′-TCCGTAGGATTCAACTGCTG-3′ |
| Gene | Primers | |
| VDAC/ Porin | Forward | 5'-ATCTGAAGACTAAGACCTCGTCG-3' |
| | Reverse | 5'-AGACCTTTCCAGACTCCTGGT-3' |
| IP3R/ Itpr | Forward | 5'- ATGGGCGACAATATAATTGGCTC-3' |
| | Reverse | 5'-GTGCTCAAGAAACCGCAAACG-3' |
| GRP75/Hsc70-5 | Forward | 5'-CGCGTACCCAAGTTTCTGC-3' |
| | Reverse | 5'-CGGAACATGCTAGAAGCTCC-3' |
| FGF21/ bln | Forward | 5'- TGTCGCCCGCTGACAATAAT-3' |
| | Reverse | 5' TTGCTGATGGGCGTGTTACT-3' |
| Ire1/ Ire1 | Forward | 5'-ATGGTAAGGAGGGCGAGCAG-3' |
| | Reverse | 5'-ATGACCGTGTACTGAGTC-3' |
| Bip(grp78)/ Hsc 70-3 | Forward | 5'- GAATCAGTTGACCACCAATCCC-3' |
| | Reverse | 5'-AACTTGATGTCGTGTTGCACA-3' |
| ATF6/Atf6 | Forward | 5'-TGAGCCTAATTCGTCTCCCAC-3' |
| | Reverse | 5'-TAGACCGCCTCTTCGTTAGAA-3' |
